# Computational Modeling and Analysis of the TGF-β-induced ERK and SMAD Pathways

**DOI:** 10.1101/2024.11.07.622480

**Authors:** Seung-Wook Chung, Carlton R. Cooper, Mary C. Farach-Carson, Babatunde A. Ogunnaike

**Affiliations:** Department of Chemical & Biomolecular Engineering; University of Delaware, Newark DE, U.S.A; Departments of Biological Sciences & Black American Studies, University of Delaware, Newark DE, U.S.A; Departments of BioSciences and Bioengineering, Rice University, Houston, TX, U.S.A

**Keywords:** TGF-β, Signal transduction, SMAD pathway, ERK/MAPK pathway, cancer

## Abstract

TGF-β, an important cytokine that plays a key role in many diseases regulates a wide array of cellular and physiologic processes via several TGF-β-driven signaling cascades, including the SMAD and non-SMAD-driven pathways. However, the detailed mechanisms by which TGF-β induces such diverse responses remain poorly understood. In particular, compared to the SMAD-dependent pathway, SMAD-independent pathways such as the ERK/MAPK pathway, which is critical in cancer progression, are less characterized. Here, we develop an integrated mechanistic model of the TGF-β-triggered ERK activation pathway and its crosstalk with the SMAD pathway, an analysis of which demonstrates how SMAD dynamics can be significantly modulated and regulated by the ERK pathway. In particular, SMAD-mediated transcription can be altered and delayed due to expedited phosphorylation of the linker of SMAD by TGF-β-activated ERK; and enhanced ERK activity, but attenuated SMAD activity, can be achieved simultaneously by fast turnover of TGF-β receptors via lipid-rafts. Also, *in silico* mutations of the TGF-β pathways reveal that the dynamic characteristics of both SMAD and ERK signaling may change significantly during cancer development. Specifically, normal cells may exhibit enhanced and sustained SMAD signaling with transient ERK activation, whereas cancerous cells may produce elevated and prolonged ERK signaling with enervated SMAD activation. These distinctive differences between normal and cancerous signaling behavior provide clues concerning, and potential explanations for, the seemingly contradictory roles played by TGF-β during cancer progression. We demonstrate how crosstalk among various branch pathways of TGF-β can influence overall cellular behavior. Based on model analysis, we hypothesize that aberrant molecular alterations drive changes in the intensity and duration of SMAD and ERK signaling during cancer progression and ultimately lead to an imbalance between the SMAD and ERK pathways in favor of tumor promotion. Thus, to treat cancer patients with a genetic signature of oncogenic Ras effectively may require *at least* a combination therapy to restore *both* the expression of TGF-β receptors and the GTPase activity of Ras.

## Background

Intricate cell-cell and cell-environment communication systems are fundamental to the proper function of individual cells in the context of a whole organism. In particular, basic cellular functions such as cell proliferation, differentiation, migration, and death, are maintained under the tight control of a dense network of secreted protein signals such as cytokines, growth factors or hormones. Among these signals, transforming growth factor-β (TGF-β) is particularly prominent because it regulates a wide variety of essential functions of virtually all human cell types, and its disruption often leads to several major diseases, including cancer [1].

TGF-β exerts potent tumor-suppressive effects on normal or pre-malignant epithelia by promoting such processes as cell cycle arrest and programmed cell death (or apoptosis). However, paradoxically, TGF-β also stimulates other physiologic processes that cancer cells exploit to their advantage, such as epithelial-mesenchymal transition (EMT), cell invasion, microenvironment modification. Consequently, in advanced cancers, TGF-β appears to play a tumor-supporting role [1]. Our current understanding of the mechanistic basis of TGF-β’s diverse but apparently contradictory roles during cancer progression remains incomplete.

What is known is that TGF-β’s versatility is associated with a variety of affiliated intracellular components. Among these, the signaling cascade consisting of SMAD proteins is relatively well characterized. This important signaling cascade begins when bioactive TGF-β binds to and brings together type I and type II TGF-β receptor serine/threonine kinases on the cell surface, whereby the activated type II receptor transphosphorylates and activates the type I receptor. The activated type I receptor, in turn, propagates the signal through phosphorylation of receptor-bound (R-) SMAD transcription factors (i.e., SMAD2/3 or SMAD1/5/8) at their C-terminal SSXS motif. The activated R-SMADs then form a heteromeric complex with co-SMAD (or SMAD4), and rapidly translocate into the nucleus where they undergo continuous nucleocytoplasmic shuttling by interacting with the nuclear pore complex. Once in the nucleus, activated SMAD complexes bind to specific promoters and ultimately regulate expression of target genes through interactions with other transcriptional co-activators and co-repressors, generating a variety of cellular responses in a cell- and context-specific manner.

A growing body of biochemical evidence now has revealed that SMAD-independent pathways that are activated by TGF-β in parallel with the canonical SMAD pathway also contribute to the diversity of TGF-β-induced responses [2, 3]. Non-SMAD signal transducers not only interact with the SMAD proteins to modulate (i.e., potentiate, synergize with, or antagonize) the SMAD pathway, but also serve as critical nodes for crosstalk with other major signaling pathways, such as receptor tyrosine kinase (RTK) pathways. One of the non-SMAD pathways that continues to attract attention is the TGF-β-activated extracellular signal-regulated kinase (ERK) mitogen-activated protein kinase (MAPK) pathway.

TGF-β-induced ERK activation clearly plays a critical role in many cellular responses in a cell-specific fashion. First, ERK can interact directly with SMAD proteins by phosphorylating the linker segments of several SMADs (including SMAD1, SMAD2 and SMAD3) to regulate ligand-induced nuclear translocation of receptor-activated SMADs [4, 5]. This negative regulation of SMADs by ERK is reported to be significant especially in oncogenic settings such as Ras hyperactivation [4]. Also, substrates of ERK (e.g., AP-1 family members) can interact with SMADs to regulate gene expression [6]. Furthermore, ERK activation is necessary for disassembly of cell adherens junctions and induction of cell motility, as part of the TGF-β-induced EMT program [7]. This TGF-β-mediated program is important not only in normal physiological processes (e.g., embryogenesis, wound healing), but also in pathological ones such as cancer metastasis. In addition, a growing number of studies have observed a high correlation between TGF-β and ERK-mediated responses. Nevertheless, our understanding of the molecular details underlying how TGF-β activates ERK and its implication for cellular behavior currently remains incomplete.

Conventionally, ERK activation is reported to be initiated by ligand-activated receptor tyrosine kinases (RTKs), typically via the Grb2/Sos complex that triggers the Ras➔Raf➔MEK➔ERK cascade. On the other hand, TGF-β signaling is initiated by its cognate receptors that possess receptor serine/threonine kinase (RSTK) activity. This raises an important question: how can such different, seemingly parallel receptor kinase (RTK vs RSTK) signaling routes be activated presumably simultaneously like a single, consolidated pathway in response to an RSTK-dedicated ligand, TGF-β, even without RTK stimuli? This question was partially answered by Lee et al. demonstrating that TGF-β receptors are dual-specificity kinases [8]. Specifically, TGF-β-activated TβRI can recruit and directly phosphorylate ShcA on tyrosine and serine residues, thus promoting the formation of a ShcA/Grb2/Sos complex. This provides a plausible molecular basis for linking the TGF-β receptor module to the downstream ERK MAPK pathway.

While such experimental evidence as this improves *qualitative* understanding of aspects of the TGF-β-induced ERK signaling cascade, there remains a significant lack of comprehensive and *quantitative* understanding of the dynamics of TGF-β signaling pathways encompassing both SMAD and ERK cascades and their interactions. To the best of our knowledge, no comprehensive mathematical model of non-SMAD signaling pathways exists in the open literature (except for our previous public, non-archival presentation [9]). In that case, what we present in this paper is the first mechanistic model of the TGF-β-triggered ERK activation pathway and its crosstalk with the canonical SMAD pathway. (While some mathematical models of TGF-β signaling [10, 11] have previously been used to investigate signal “crosstalk”, it is important to note that these studies are concerned with “crosstalk” between SMAD pathways induced by *different* types of TGF-β superfamily receptors. By contrast, our current study is concerned with “crosstalk” of an entirely different nature, namely, the interplay between distinct and seemingly unrelated pathways, SMAD and ERK signaling pathways, induced by the *same* TGF-β receptor types (TβRI and TβRII)).

Our model is used to develop quantitative insight into how crosstalk among the various TGF-β branch pathways influences overall system behavior. The model also is used to simulate the dynamic behavior of cancerous systems, from which we generate hypotheses regarding potential mechanisms for how TGF-β’s tumor-suppressive roles may seem to morph into tumor-promoting roles. Finally, we discuss potential therapeutic implications for treating cancers of epithelial origin that are TGF-β signaling-related.

## Methods

### Model Formulation

To construct a more comprehensive TGF-β network signaling model, we extended our previously published model of the SMAD pathway [12] by incorporating TGF-β-triggered ERK activation mechanisms. In particular, we postulated for the ERK pathway module, several essential molecular reactions based on up-to-date experimental findings reported in the literature, and adapted existing, well-cited mathematical models of ERK signaling such as the Schoeberl model [13]. We describe below the essential molecular processes on which the model structure is based. We do not describe in detail those biochemical processes featured in existing models that we simply adapted for our use; rather, we focus our detailed descriptions on newly proposed mechanisms. Also, while the incorporation of additional possible mechanisms including a variety of feedback loops may be of interest for future studies, such mechanisms are not of direct relevance to this study and hence lie outside of the intended scope.

### Receptor activation

The active form of TGF-β ligand binds to the extracellular domain of dimeric type II receptor (denoted as RII in this study) and forms a catalytically active TGFβ-RII complex. The activated TGFβ-RII complex then engages with and activates type I receptor (denoted as RI), forming a TGFβ-RII-RI oligomer at the cell surface, which is able to trigger downstream signaling.

### Different endocytic routes of TGF-**β** receptors

It is well documented that TGF-β receptors are internalized via two major endocytic routes: clathrin-mediated and lipid raft-mediated endocytosis [14]. Unlike growth factor-activated tyrosine kinase receptors, TGF-β receptor endocytosis is not initiated or facilitated upon binding of TGF-β to the receptors; the receptors are constitutively internalized and recycled [15, 16]. Clathrin-mediated internalization of TGF-β receptors to early endosomes is known to promote SMAD-dependent signaling via scaffold proteins such as SARA, whereas cholesterol-rich lipid-raft/caveolae-mediated endocytosis leads to the degradation of TGF-β-activated receptors, therefore turning off TGF-β signaling [15].

Interestingly, lipid rafts/caveolae may serve as a subcellular location to promote SMAD7-mediated degradation of TGF-β receptors, while also participating in some of the specific TGF-β signaling pathways. For example, Zuo and Chen reported that the localization of receptors at various plasma membrane regions can determine the downstream signaling of TGF-β and thus influence cellular response outcomes [17]. They reported that although receptor hetero-complexes can be formed in both lipid raft and non-raft membrane compartments in response to TGF-β, the localization of receptors in lipid rafts, but not clathrin-coated pits, is important for TGF-β-induced MAPK activation, leading to epithelial cell plasticity. In other words, it is possible that the diversity of TGF-β responses depends on the compartment-specific localization of TGF-β receptors. Consequently, we postulate that clathrin-dependent receptor internalization to early endosomes promotes SMAD-mediated signaling, whereas lipid-raft-dependent endocytosis is dedicated to non-SMAD signaling including ERK signaling, as well as to the degradation of ligand-activated receptors, leading to the attenuation or termination of SMAD signaling.

### ShcA activation by TGF-β receptors

As noted earlier, Lee and colleagues demonstrated for the first time that in response to TGF-β, activated type I receptor recruits and directly phosphorylates ShcA proteins on tyrosine and serine [8], providing a basis for one possible mechanism of TGF-β-triggered ERK activation. Once activated, ShcA can associate with Grb2 and Sos proteins, thereby activating the well-characterized ERK MAPK cascade. Based on this, we postulate that ligand-activated receptor complex internalized via lipid-raft associates directly with and activates ShcA, initiating ShcA/Grb2/Sos-mediated downstream ERK signaling.

### The Ras/Raf/MEK/ERK cascades

For the downstream signaling layout of the MAPK pathway, specifically from ShcA activation to double-phosphorylated ERK (denoted as ppERK), we adapted previously reported mathematical models for the EGF signaling pathway, in particular the Schoeberl model [13] as follows: phosphorylation of Shc protein by ligand-activated receptors leads to the formation of the Shc-Grb2-Sos complex, which activates Ras protein, a member of the GTPase family, by catalyzing the exchange of Ras-bound GDP with GTP. Once in its GTP-bound state, Ras can associate with and activate MAP kinase (MAPKKK), Raf. The activated Raf then phosphorylates and activates MAP kinase (MAPKK), MEK1 and MEK2 (consolidated and denoted as MEK in our model), which in turn phosphorylate and activate MAP kinase (MAPK), ERK1 and ERK2 (consolidated and denoted as ERK in our model).

### SMAD phosphorylation and complex formation

The mechanism of how R-SMAD is activated and forms a complex with co-SMAD is summarized in our previous paper [12]. Briefly, ligand-activated receptor complex that is internalized into early endosomes associates with and phosphorylates R-SMAD. Once phosphorylated, R-SMAD forms a complex with SMAD4 and participates in transcription of target genes. Because the current model focuses exclusively on cytoplasmic events, other molecular mechanisms occurring in the nucleus are not considered in this study.

### ERK-induced phosphorylation of SMAD

Although C-terminal SXS phosphorylation by ligand-activated type I receptor is the key event in SMAD activation, additional phosphorylation by intracellular protein kinases can also regulate SMADs positively and negatively. In particular, the regions that join two conserved polypeptide segments (e.g., the MH1 and MH2 domains) of SMAD (or the so-called linker region) are known to be phosphorylated by MAPKs including ERK. The linker phosphorylation gives rise to nuclear exclusion of SMAD proteins by blocking SMAD interactions with the nuclear pore complex, or by increasing the affinity of SMADs for a cytoplasmic anchor or a nuclear export molecule [4, 18]. This linker phosphorylation does not seem to prevent receptor-mediated SMAD phosphorylation [5, 19]. Therefore, we assume that ligand-induced ERK directly phosphorylates the linker region of both receptor-phosphorylated and unphosphorylated R-SMAD proteins, and that the linker-phosphorylated SMAD also can be phosphorylated in its SXS motif by ligand-activated receptor complex.

### SMAD dephosphorylation

Like phosphorylation, SMAD dephosphorylation is a critical event in TGF-β signaling because it controls and terminates TGF-β signaling in order to maintain tissue homeostasis and normal cellular responses. Both receptor-phosphorylated C-terminal SXS motif of SMAD and ERK-phosphorylated linker region of SMAD are dephosphorylated by specific phosphatases. Several SMAD-specific phosphatases have been reported in the literature. PPM1A was identified as a SMAD2/3 SXS-motif specific phosphatase [20], whereas SCP1/2/3 were found to dephosphorylate specifically SMAD2/3 linker [21] *in vivo* as well as *in vitro*. Although these phosphatases are found to be localized primarily in the nucleus, R-SMAD dephosphorylation cannot be considered as an exclusively nuclear event. This is because C-terminal dephosphorylation of ligand-activated SMAD can also be mediated by non-nuclear phosphatases such as pyruvate dehydrogenase phosphatase (PDP) [22, 23]. Because this study (and the resulting model) deals with cytoplasmic signaling events only, for simplicity, we consider as the signal turn-off mechanisms cytoplasmic dephosphorylation of both C-terminus- and linker-phosphorylated SMAD.

### Sos phosphorylation by ppERK

Activated ERK is known to feed back to the pathway activation process at several points [24]. In particular, activated ERK inactivates Sos by direct phosphorylation, which triggers the disassembly of Grb2-Sos complex, thereby strongly inhibiting Sos-dependent Ras activation [25–28]. Consequently, we assume that double-phosphorylated ERK (i.e., ppERK) phosphorylates all Sos-containing species and dissociates Sos from any of its complex.

### Model Overview

A simplified schematic representation of the model structure is shown in Figure 1 (See Figure S1 in Additional File 2 for the biochemical reactions and other details regarding the model). The input to the model is TGF-β concentration (shown in blue in Figure S1); the responses of interest are phosphorylated RSmad-Smad4 complex (green in Figure S1) and double phosphorylated ERK (red in Figure S1). Based on two well-mixed compartments—the extracellular and the cytoplasmic compartments—the resulting model is a system of 61 non-linear ordinary differential equations (ODEs), with 121 kinetic parameters arising from chemical reactions represented by mass action kinetics and Michaelis-Menten kinetics (See Table S1 in Additional File 1 for the complete set of equations.) The cell is idealized as a sphere with a cell volume of 1.5 pL and a cytoplasmic volume chosen to be three quarters of the total cellular volume [29], not unrealistic for a depolarized cancer cell. Simulations are generated using the ‘*ode15s*’ routine of MATLAB 7.1 (The MathWorks, Inc.) to integrate the model equations.

**Figure 1.**
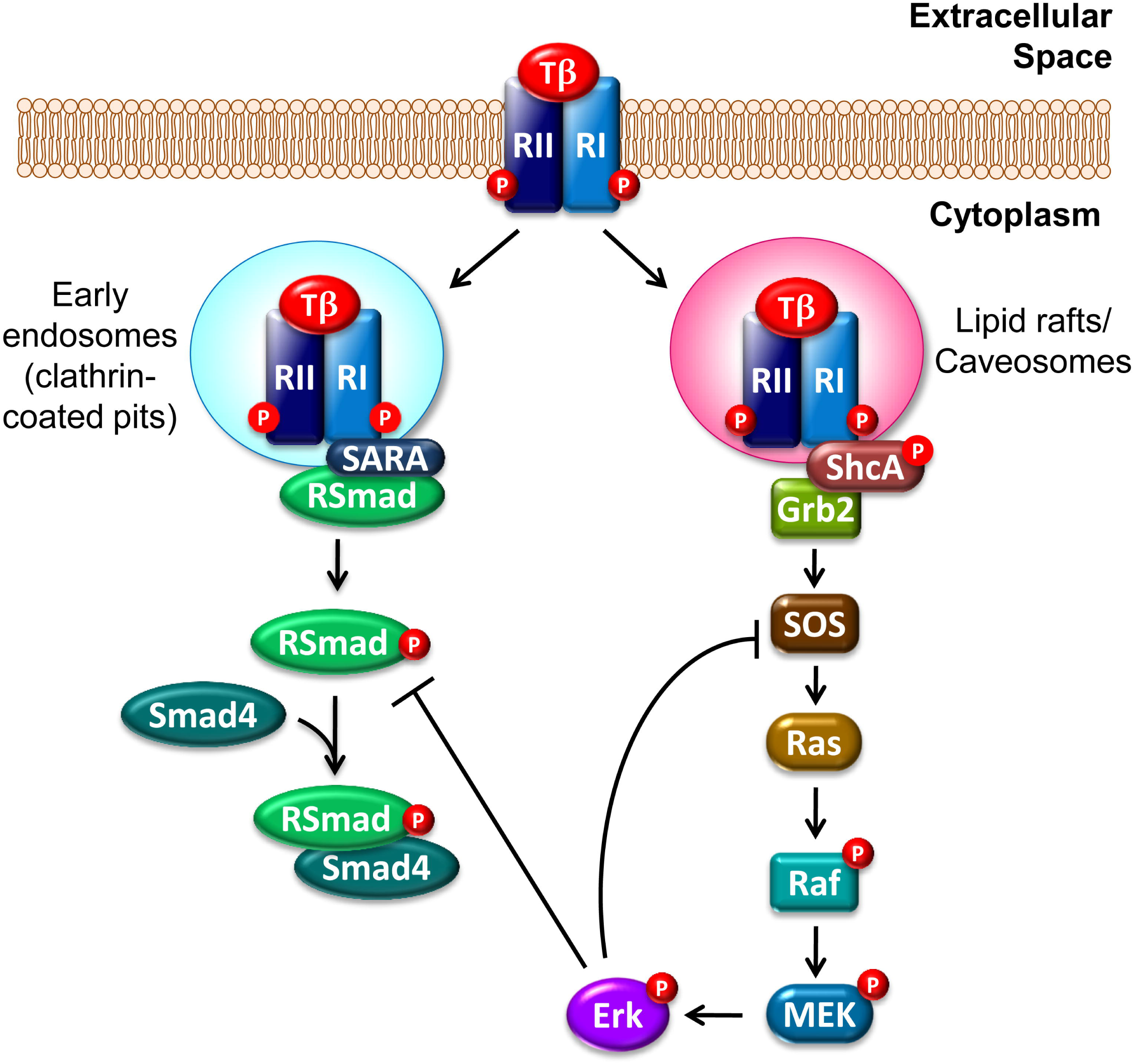
Simplified schematic of the TGF-β-induced SMAD and ERK signaling pathways.

### Initial Conditions

As stated earlier, various aspects of our model mechanisms were adapted from existing mathematical models of individual ERK and SMAD signaling pathways, including our own group’s models [12, 13, 30–32]. As such, the pre-stimulus steady-state values for species with non-zero initial conditions (i.e., TGF-β receptors, R-SMAD, SMAD4, Shc, Grb, Sos, RasGDP, Raf, MEK, ERK, and phosphatases) were obtained from these studies. In particular, for the basal levels of TGF-β receptors, we assumed that 15% of total receptors (5,000 copies per cell for each receptor type) are present at the cell surface and the remaining 85% of total receptors are constitutively internalized either to early endosomes or to lipid-rafts in approximately equal amounts [12, 32]. The basal levels of all other species not listed above are assumed to be zero initially (See Table S2 in Additional File 1).

### Model Parameter Estimation

One of the most difficult challenges of effective dynamic modeling of cellular signal transduction systems is parameter estimation. This is because, to achieve any reasonable degree of fidelity, the models almost always consist of a relatively large number of model parameters, but only a paltry amount of relevant data is available for estimating unknown values. Furthermore, much of the available data are almost always obtained from experiments that were not designed specifically for parameter estimation [33]. Parameter estimation for signal transduction models must therefore be carried out with care. Our approach is summarized as follows (also discussed in our previous study [12]):

1. *Initial Rough Estimation*: Initial values for many parameters were obtained directly from previous models, primarily our SMAD pathway model [12] and the Schoeberl models [13]. Several kinetic parameter values were determined through extensive literature search; others were computed from available *in vitro* experimental data. The remaining unknown parameters were assigned initial estimates and reasonable upper and lower bounds by comparison with similar circumstances in the literature (e.g., similar components in other published signaling pathway models) and from known physical limitations (e.g., diffusion-limited rates). See Table S3 in Additional File 1 for additional specific details.

2. *Parametric Sensitivity Analysis*: To determine parameters that are the most sensitive and therefore need more precise estimation, we used the set of initial estimates determined in Step 1 to carry out local parameter sensitivity analysis in order to quantify the effect of parametric changes on the four system responses of interest (see Figure 2) for which experimental data are available. The calculations are based on the following expression for the normalized sensitivity coefficient (NSC):

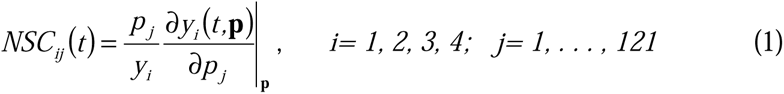

**Figure 2.**
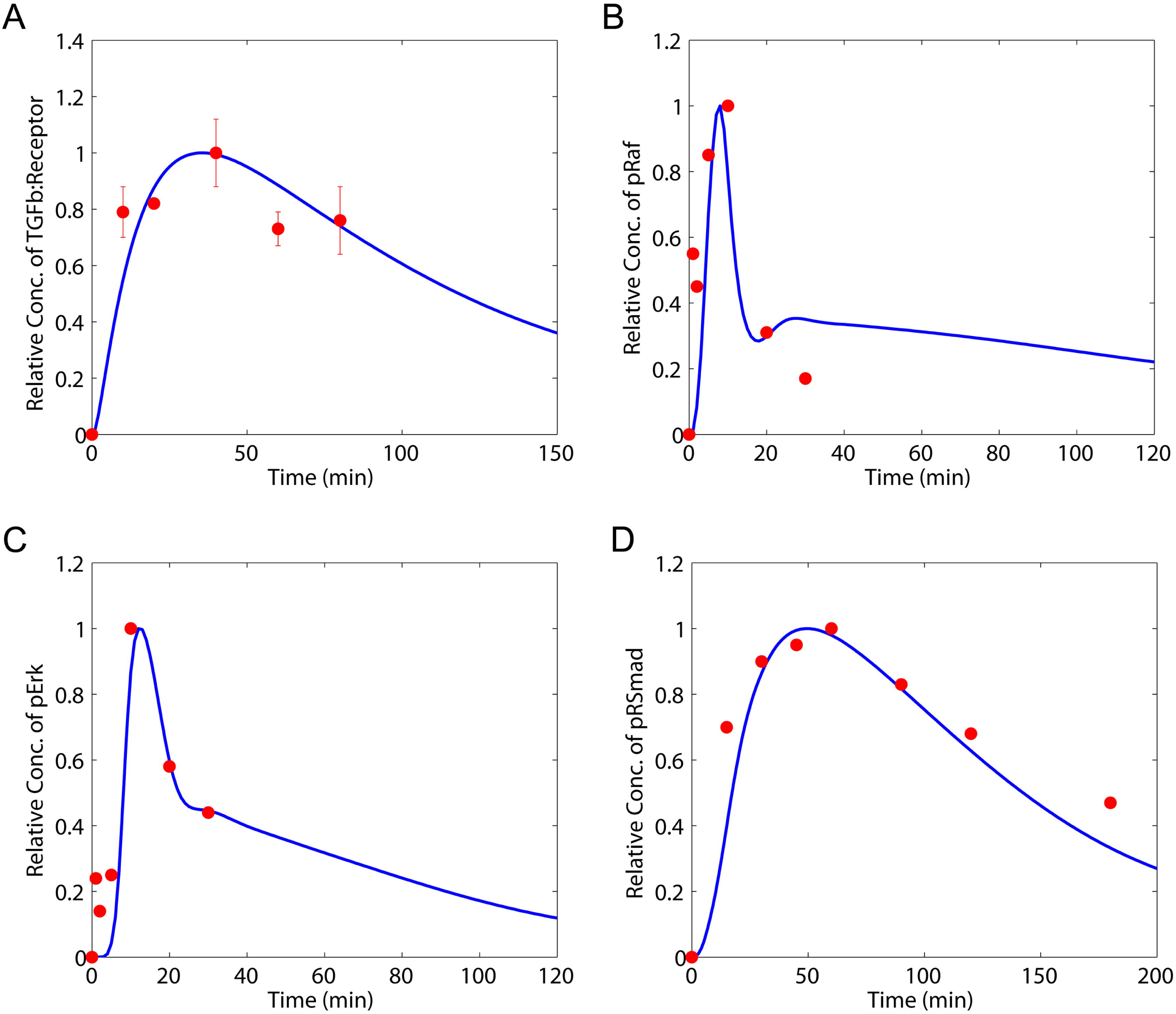
Model fit to experimental data. Model prediction (solid line) vs data (filled circle) for (A) TGF-β-bound receptors in early endosomes [34] ; (B) phosphorylated Raf [8]; (C) phosphorylated ERK [8]; and (D) phosphorylated SMAD2 [35].

where *y_i_* represents the system response variable in question, and **p** denotes the vector of kinetic parameters. A total of 24 parameters (listed in Table S3) were selected for more precise estimation because of their high NSCs and/or because we have little or no confidence in their initial values.

3. *Model Calibration*: We calibrated our model by fitting our model predictions *simultaneously* to published *in vitro* experimental time course data on: (i) TGF-β-bound receptors in early endosomes [34]; (ii) phosphorylated Raf [8]; (iii) phosphorylated ERK [8]; and (iv) phosphorylated SMAD2 [35], obtained from the indicated publications. The raw data, available in the form of immunoblot image, were quantified with the ‘Image Processing Toolbox’ in MATLAB 7.1, normalized using the largest value of each data set, and compared to the corresponding similarly normalized model predictions. The model fit was carried out using MATLAB’s non-linear least squares solver, ‘*lsqnonlin*’.

4. *Identifiability*: To identify whether the unknown model parameters can be uniquely estimated from the available data, a “practical identifiability” analysis was performed, following our previous studies [33, 30, 12]. For this current study, tolerances were generously set at ± ∼50% and the results are summarized in Table S3.

5. *Identifiable Parameter Estimate Refinement*: Estimates for identifiable parameters were further refined by repeating Step 3 (least squares estimation) and Step 4 (local identifiability test) until the “best” estimates of this subset of parameters were determined.

The final result is shown in Table S2 in Additional File 1.

## Results and Discussion

Unless stated otherwise, each simulation represents system response to a single step of magnitude 80 pM in TGF-β, introduced at time *t=0*.

### Model and Parameter Fit Evaluation

A direct comparison between the normalized *in vitro* experimental data and the corresponding optimized model prediction is shown in Figure 2. Considering that the model was fit to these four different data sets *simultaneously*, the resulting agreement between model prediction and data is very good. The implications are twofold: (i) parameter estimates determined by the optimization exercise are fairly representative of the true (but unknown) model parameter values; (ii) the proposed model structure itself is a very reasonable mathematical representation of a far more complex reality. The inevitable discrepancies between model predictions and experimental data are attributable to multiple factors, including simultaneous fitting to experimental data generated under non-identical conditions; potential lack of direct correlation between *in vitro* and *in vivo* measurements; model non-linearity causing multiple local minima in parameter estimation, etc.; all of which were discussed in detail in our previous study [12]. Taking all these factors into consideration, we conclude that the model captures the dynamics of TGF-β signaling quite well.

### Independent Model Validation

Before proceeding to use the model, it is important to validate its predictions against a *different* set of independent experimental data, *without further adjusting any model parameter*. To test the model’s ability to capture other aspects of the system dynamics under different experimental conditions, we compared the model predictions to a different collection of four independent experimental data sets obtained from the literature: (i) phosphorylated ShcA [8]; (ii) GTP bound Ras [36]; (iii) phosphorylated SMAD2 in response to a step input of TGF-β [15]; and (iv) phosphorylated SMAD3, in response to a 30 min pulse input of TGF-β [20]. Figure 3 shows a comparison of the model predictions *directly* to corresponding four separate and independent experimental data sets. Note that the model parameters were not modified in order to obtain the indicated fit. Even under these stringent conditions the model still shows reasonable agreement with the dynamics of a set of observables totally different from those used for model fitting. We conclude therefore that the model provides a very good representation of the dynamic behavior of TGF-β-induced simultaneous activation of *both* the SMAD and ERK pathways.

**Figure 3.**
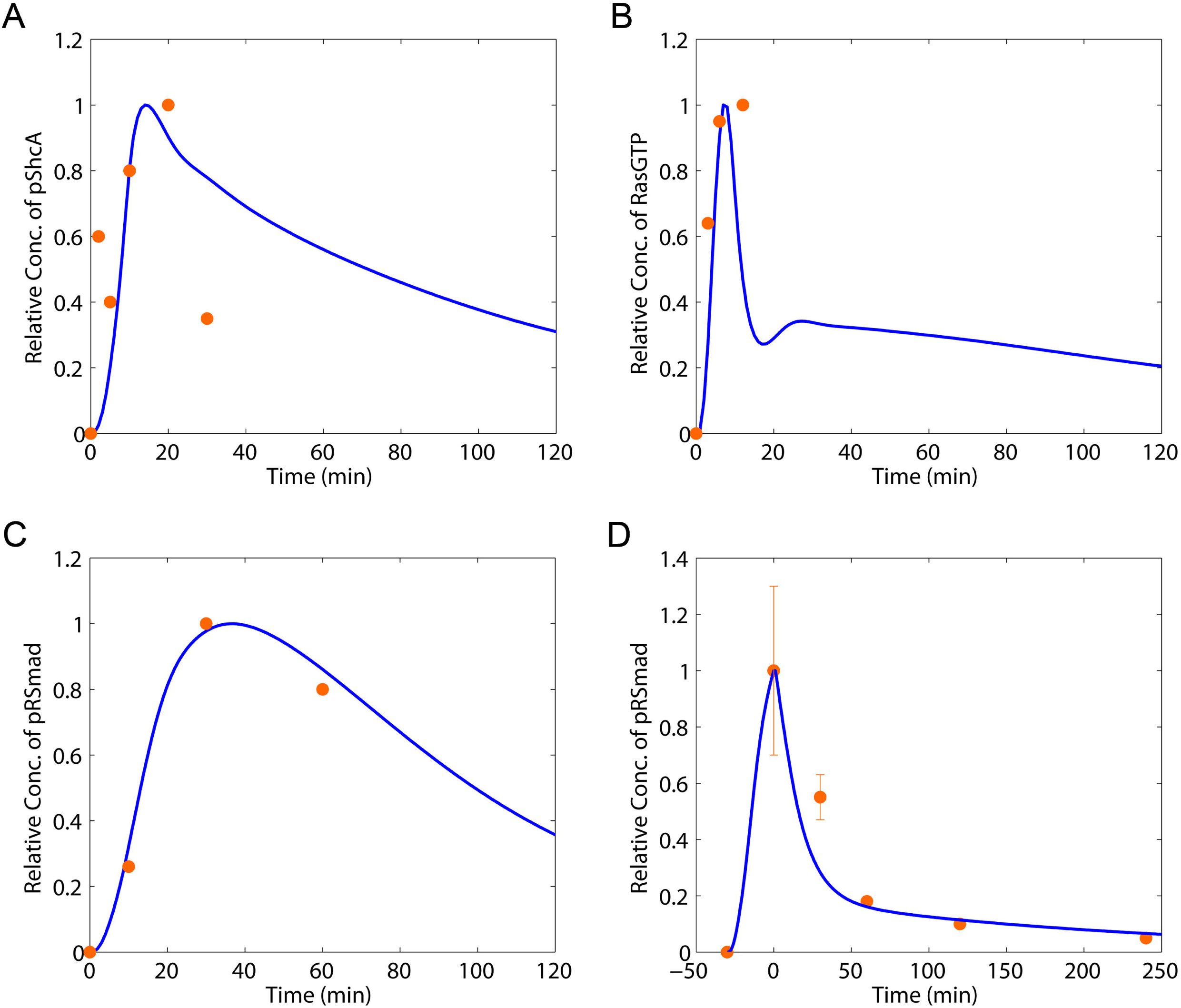
Model validation with independent experimental data. Model prediction (solid line) versus data (filled circle) for (A) phosphorylated ShcA [8]; (B) GTP bound Ras [36]; (C) phosphorylated SMAD2 in response to a step input of TGF-β [15]; and (D) phosphorylated SMAD3 in response to a 30 min-pulse input of TGF-β [20].

### Parametric Sensitivity Analysis

To understand quantitatively which aspects of the signaling network most influence system behavior, we carried out parametric sensitivity analysis for the two primary outputs of interest: double-phosphorylated ERK (ppERK) and phosphorylated RSMAD-SMAD4 complex (pRSMAD-SMAD4). First, Figure 4A shows normalized sensitivity coefficients (computed using Eq. 1) for ppERK, as a function of time, for the 10 most important parameters (parameters for which the maximum normalized sensitivity coefficient exceeds 2.2 in absolute value at any point in time). These important parameters are seen to fall naturally into two groups based on their temporal profiles. Group 1 parameters affect the output variable strongly immediately after ligand stimulation, but their influence decreases rapidly thereafter. Parameters in this group are involved in the following reactions: binding between ligand and TβRII (*p_1_*); raft-mediated internalization of receptor complex (*p_5_*); and complex formation of pRc-pShc-Grb-Sos (*p_11_*). On the other hand, the effect of Group 2 parameters on the output variable is itself quite dynamic, with normalized sensitivity coefficients that change significantly in the short term (within the first 30 mins), which subsequently increase (in absolute value) monotonically over time. The parameters in this group are associated with the following reactions: complex formation of pRc-pShc-Grb-Sos (*p_27_*); Ras-GTP activation (*p_30_*, *p_32_*); Raf activation (*p_33_*); Ras-GTP deactivation (*p_36_, p_37_*), and pRaf deactivation (*p_38_*).

**Figure 4.**
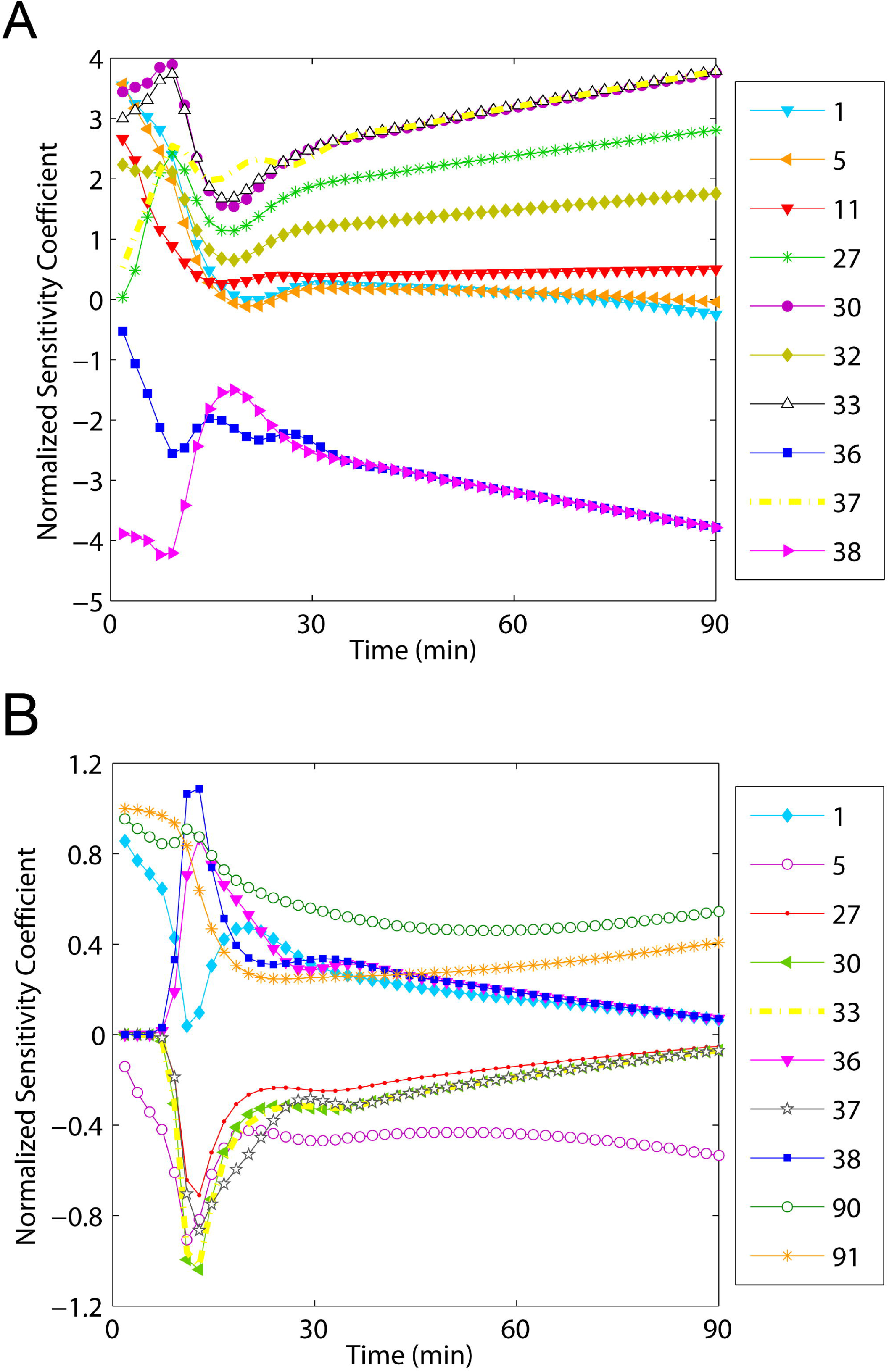
Model parameter sensitivities for select parameters with the greatest influence on key outputs. (A) ppERK and (B) pRSMAD-SMAD4 complex. Numbers in each legend refer to parameter indices shown in Table S3.

Interestingly, the point in time at which the dynamics of the two groups differ most markedly is associated with the formation of the receptor-bound Sos complex, which then activates Ras protein, followed by the downstream MAPK cascade. Indeed, Group 1 parameters are related to the initial phase of ERK activation at the receptor level, whereas Group 2 parameters are associated with the downstream ERK activation cascade. In particular, the ups and downs of the sensitivities of Group 2 parameters correlate with the dynamic changes associated with the activation of Raf, MEK, and ERK in the hierarchical MAPK phosphorylation cascade. ***These results highlight the importance of Ras protein as a critical regulatory node that couples the upstream receptor module with the downstream MAPK cascade module.*** The significance of this insight will become evident subsequently.

For the pRSMAD-SMAD4 complex, Figure 4B shows that the 10 most important parameters (with maximum normalized sensitivity coefficient exceeding 0.6 in absolute value at any point in time) are associated with the following reactions, in order of the parameter indices: (i) association between ligand and TβRII (*p_1_*); (ii) Raft-mediated internalization of ligand-activated receptor complex (*p_5_*); (iii) formation of pRc-pShc-Grb-Sos complex (*p_27_*); (iv) binding of Ras-GTP to receptor complex (*p_30_*); (v) phosphorylation of Raf (*p_33_*); (vi, vii) Ras-GTP deactivation (*p_36_*, *p_37_*); (viii) dephosphorylation of pRaf (*p_38_*); (ix) clathrin-mediated internalization of ligand activated receptor (*p_89_*); and (x) binding of RSMAD to receptor complex in early endosomes (*p_90_*). These results show that except for two parameters associated with the SMAD pathway (*p_89_* and *p_90_*), the remaining parameters change significantly at approximately 10 min, which correlates with the approximate peak time of ppERK response. Indeed, many of the latter set of parameters overlap with the parameters associated with ERK activation. ***Collectively, this indicates clearly the importance of ERK signaling in regulating the dynamics of SMAD signaling at the cytoplasmic level***. This means that SMAD-mediated transcription and cellular responses may be affected significantly by the dynamic behavior of TGF-β-triggered ERK signaling. The obvious implication is that the SMAD pathway is strongly influenced by other signaling pathways—a conclusion that could not possibly be drawn from previous TGF-β signaling models that focused on the SMAD pathway in isolation.

Observe, however, that the mere presence of crosstalk does not necessarily mean that the implied interactions and cross-influences are all uniformly strong or consequential. By identifying the influence of non-SMAD signaling components as non-trivial, the foregoing analysis justifies the necessity of incorporating non-SMAD signaling for a more comprehensive understanding of TGF-β signaling.

### TGF-β Dose-Responses

It is well established that cellular characteristics are regulated not just by the diversity in the *type* of extracellular ligands and other signaling molecules stimulating the cell, but also by variations in the concentrations of each of these molecules. To explore how different TGF-β doses affect the system outputs, we simulated cellular responses to eight different doses of TGF-β (step changes of 4, 8, 40, 80, 400, 800, 4000, and 8000 pM), holding all other conditions constant, including initial conditions and kinetic parameters. Figure 5A shows that peak activity of pRSMAD-SMAD4 complex increases significantly as TGF-β concentration increases until saturation is reached whereby any further increase in stimuli concentration no longer generates noticeable increase in the response. The increase in TGF-β level also speeds up the overall dynamics of ligand-activated SMAD complex, reducing the time taken to reach maximum activity and steady state. The result for ppERK is similar in general (Figure 5B). However, in contrast to what is observed with the SMAD complex, whose activity in response to the even the lowest TGF-β dose (4 pM) remains observable over a simulation time of 500 min, the activity of ppERK in response to the *second* lowest TGF-β dose (8 pM) remained unnoticeable over the entire simulation time. In addition, for all concentrations of TGF-β stimuli, ERK activation is transient, while activated SMAD complex maintains its activity over the entire simulation time. In other words, the amount of TGF-β stimulus required to generate an observable response is higher for ppERK than for activated SMAD complex; furthermore, the effect of TGF-β stimuli is sustained for a much longer period of time for the activated SMAD complex. These results reveal that SMAD activation may be more sensitive to variation in the TGF-β concentration than the ERK activation. In other words, the transcription of SMAD-inducible genes is more likely to arise in response to low levels of TGF-β stimulation while ERK activation signaling is still dormant. ***These in-silico dose-response results therefore reveal how the strength of TGF-***β ***stimulus can elicit different cellular responses through different levels of activation of the key signaling components SMAD and ERK***.

**Figure 5.**
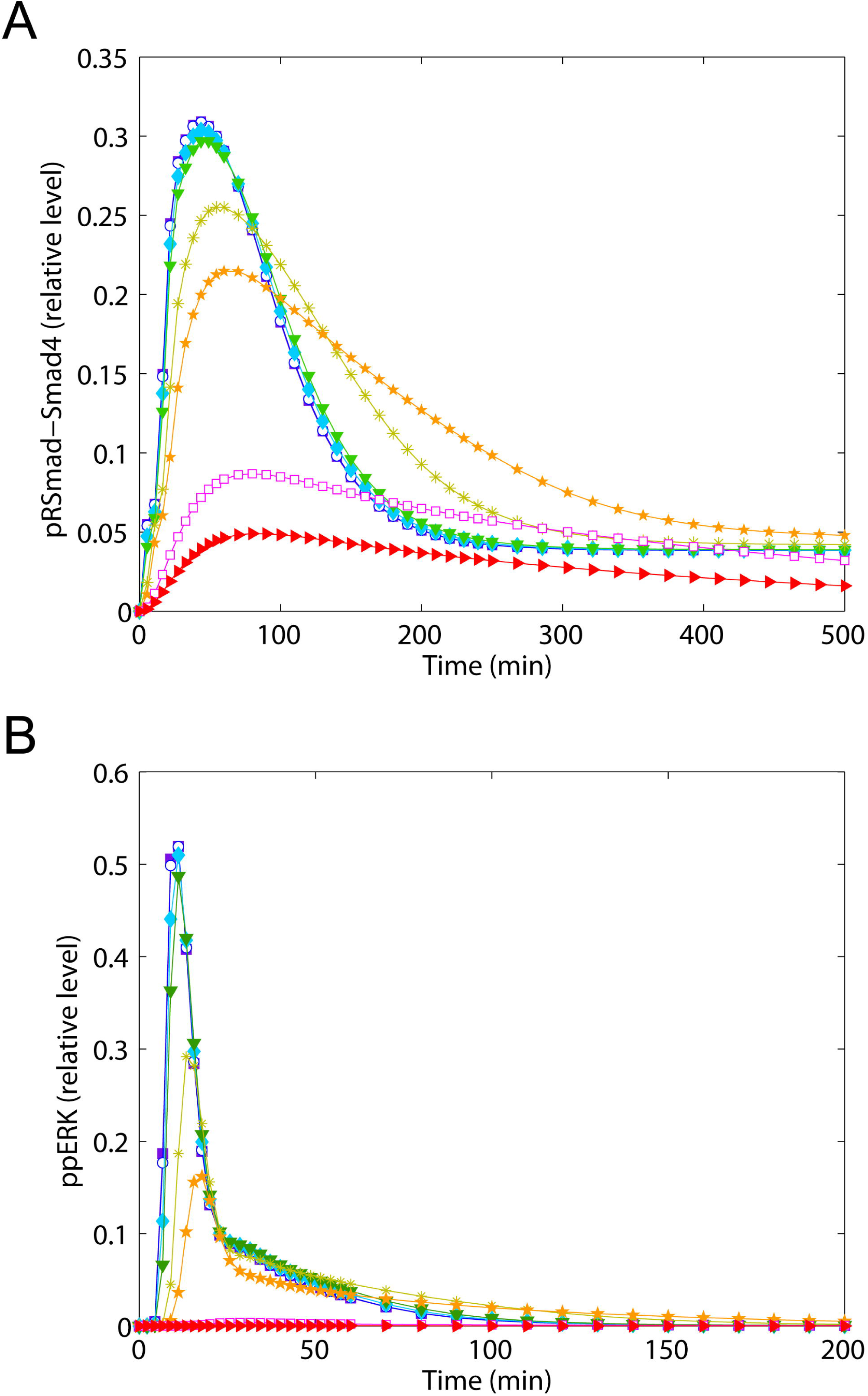
Dynamic responses to TGF-β stimuli of various concentrations. (A) pRSMAD-SMAD4 complex and (B) ppERK in response to: step changes in TGF-β concentrations of 4 (red), 8 (pink), 40 (orange), 80 (olive), 400 (green), 800 (cyan), 4000 (blue), and 8000 (violet) pM.

### Effect of Crosstalk between SMAD and ERK

Because signaling pathways rarely function in isolation, we examined the effect of interactions between SMAD and ERK signaling on the dynamics of TGF-β-mediated system responses. As noted earlier, it is known that SMAD activity can be regulated by crosstalk between SMAD and MAPKs. Specifically, ERK directly phosphorylates several sites of the linker region connecting the MH1 and MH2 domains of SMAD proteins [37]. The linker phosphorylation can lead to nuclear exclusion of and thereby attenuated transcription by ligand-activated SMADs, either by blocking SMAD interactions with the nuclear pore complex, or by increasing the affinity of SMADs for a cytoplasmic anchor or a nuclear export molecule [4, 18].

To explore how crosstalk between SMAD and ERK signaling affects TGF-β-induced responses, we simulated the system response to a single step of TGF-β, magnitude 80 pM, under the following conditions: (i) no crosstalk between SMAD and ERK; (ii) a rate of dephosphorylation of the ERK-phosphorylate linker 10 times more rapid than nominal; and (iii) a rate of phosphorylation of the ERK-phosphorylate linker 10 times more rapid than nominal. Figure 6A shows that under the first two conditions, where either crosstalk between SMAD and ERK is completely blocked (magenta), or the ERK-phosphorylated linker of SMAD is rapidly dephosphorylated (green); there is no significant increase in the receptor-activated SMAD signal compared to the nominal case (blue). On the other hand, rapid phosphorylation in the linker region of SMAD by ERK (red) leads to attenuated and sluggish dynamics of receptor-activated SMAD complex, while it increases the level of ERK-phosphorylated SMAD, which does not enter the nucleus to participate in transcription. These results imply that if crosstalk between SMAD and ERK is moderate, cells will exhibit sufficiently strong SMAD-mediated responses to TGF-β. On the other hand, if crosstalk between SMAD and ERK is much stronger (such that TGF-β-induced ERK strongly inhibits receptor-activated SMAD translocation into the nucleus), upon TGF-β stimulation, SMAD-inducible genes that are sensitive to signal intensity and/or duration may not be expressed appropriately in terms of degree and timing, while SMAD-independent genes induced via other signaling routes including ERK signaling may be strongly expressed for longer times, and potentially override SMAD-mediated responses and determine net cellular outcome.

**Figure 6.**
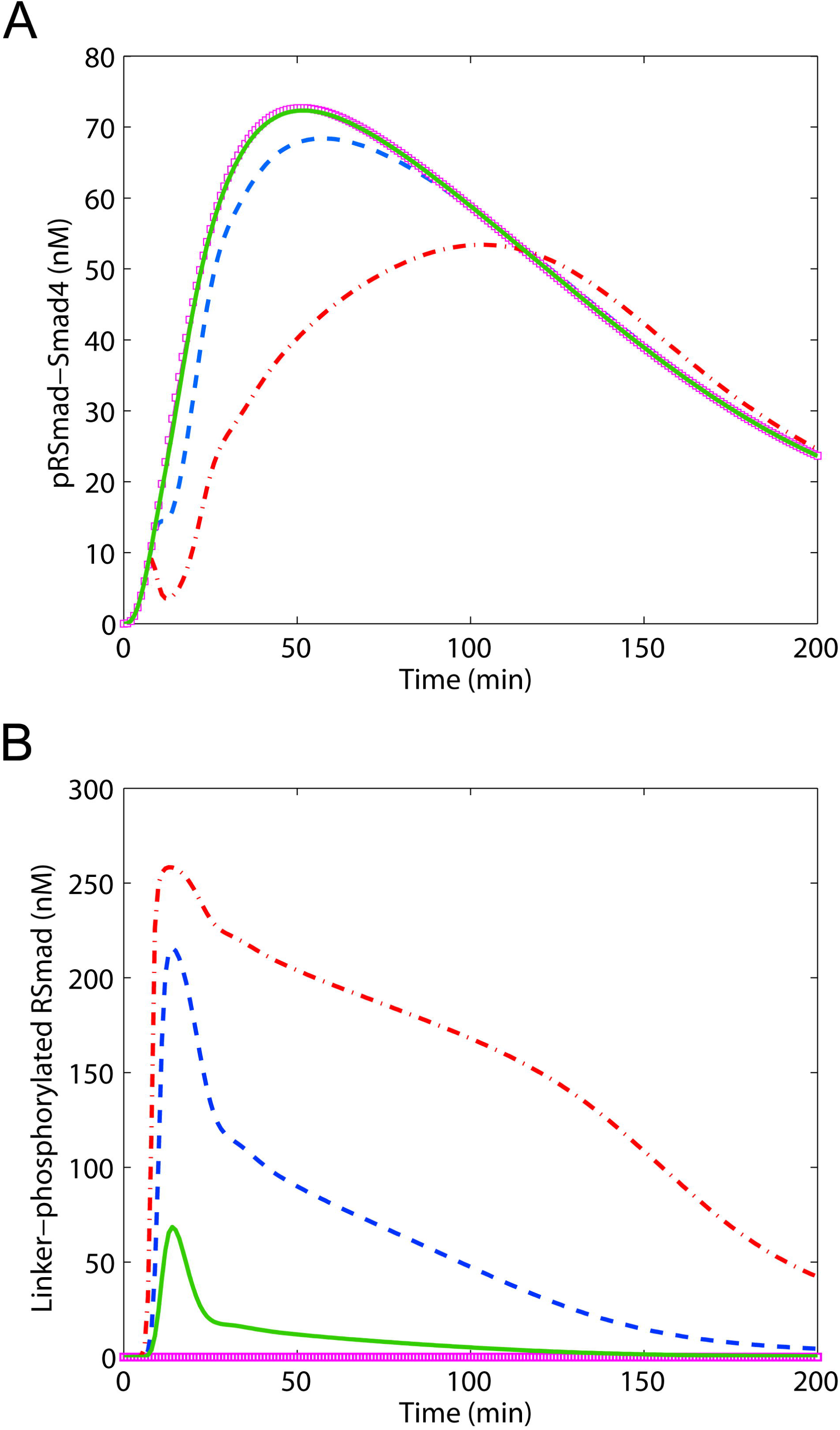
Effect of crosstalk between SMAD and ERK pathways on dynamic responses: (A) receptor-phosphorylated RSMAD-SMAD4 complex and (B) ppERK-phosphorylated RSMAD in the linker region. Rate of linker *phosphorylation* set to zero (magenta); increased 10-fold (red); rate of linker *dephosphorylation* increased 10-fold (green); nominal conditions (blue).

### Effect of Unbalanced Receptor Endocytosis

Our model assumes that different receptor endocytic routes are dedicated to specific downstream signaling cascades (i.e., lipid raft-mediated endocytosis for ERK activation, and non-lipid raft-mediated endocytosis for SMAD activation). Intuitively, therefore one would expect that any changes in these receptor endocytosis mechanisms will significantly alter observed system dynamics in predictable ways. Indeed, the foregoing parameter sensitivity analysis indicates that both receptor internalization steps are important to the response of the two system outputs of interest. One may, therefore, expect that tipping the balance between these two endocytic pathways to one side or the other will likely make one pathway’s response more dominant at the expense of the other. To test the validity of this intuitive expectation, we increased 10-fold the rate constant related to raft-mediated internalization, leaving the corresponding value related to non-raft-mediated internalization unchanged, and vice versa.

Figures 7A-C show that compared to nominal conditions (blue), when non-raft (or clathrin)-mediated internalization is predominant (red), ligand-activated SMAD signaling is enhanced and sustained (A), particularly after peak activity is attained, whereas ligand-activated ERK signaling is lowered significantly, barely registering initially and eventually appearing to turn off completely with time (C). ERK-phosphorylated SMAD (B), a surrogate of ERK activity, shows similar characteristics. The implication is that, under these conditions, the predominance of non-raft (or clathrin)-mediated internalization causes SMAD-mediated cellular outcomes to be preferentially induced, consistent with our intuitive, *à-priori* expectations. On the other hand, when raft-mediated receptor trafficking is predominant (green in Figures 7A-C), SMAD activation becomes significantly attenuated (A), as expected. However, contrary to expectations, the intensity of ligand-activated ERK signaling (C) is not enhanced significantly; instead, it is sustained for much longer than under nominal conditions. The same is true for ERK-phosphorylated SMAD (B). Why does a dominant raft-mediated route *not* lead to elevated activity of ppERK, as one would intuitively expect, and as was the case for SMAD activation under dominant non-raft internalization? This is because TGF-β receptor endocytosis to lipid raft has two distinct effects: (i) it helps the activation of ERK signaling and (ii) also promotes the degradation of the ligand-activated receptors [15, 38]. ***In other words, faster receptor internalization by lipid-rafts renders more receptors susceptible to rapid ligand-induced degradation, limiting the availability of receptors for the activation of ERK signaling*.**

**Figure 7.**
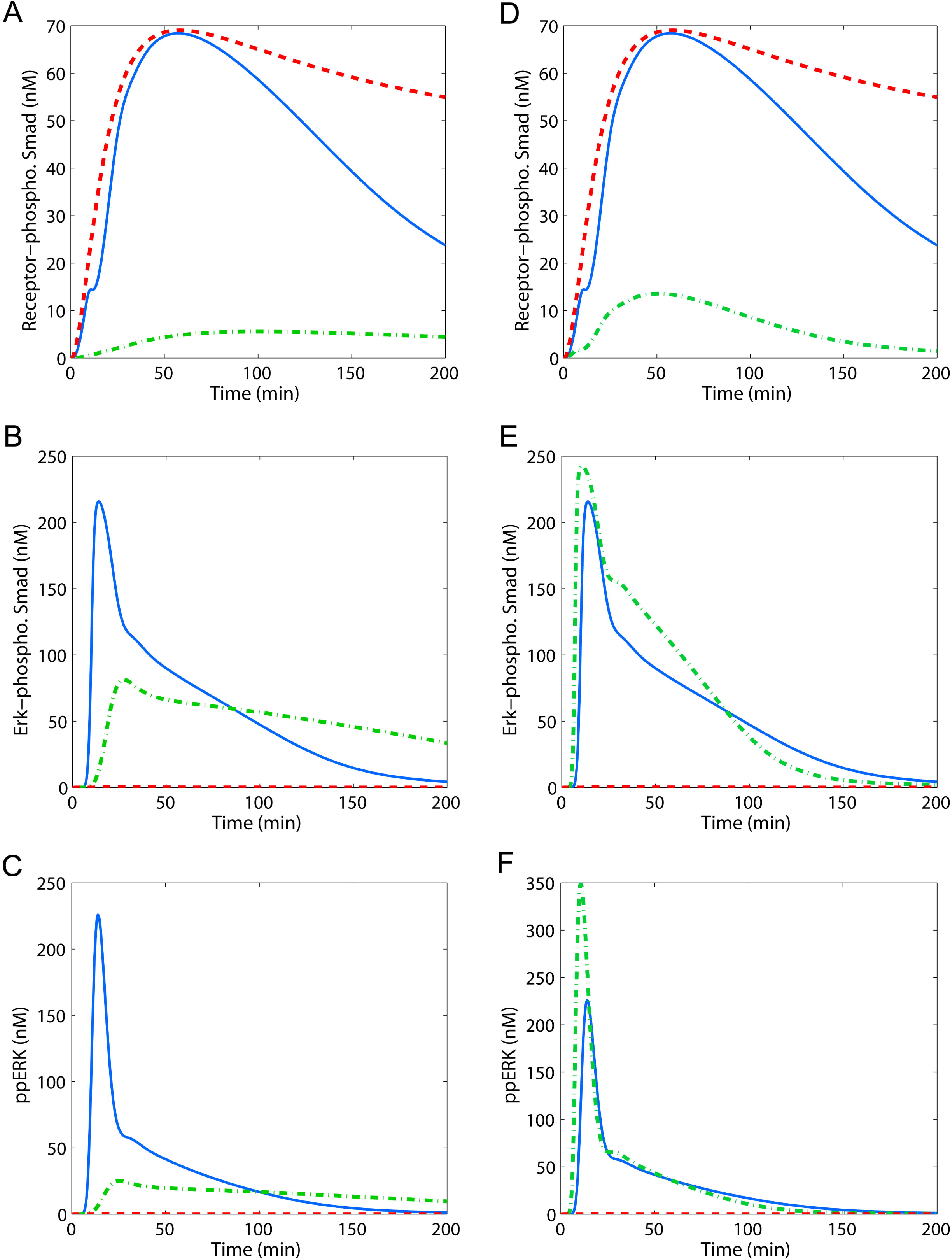
Effect of biased receptor endocytosis on key system outputs: (A and D) pRSMAD-SMAD4 complex, (B and E) ppERK-driven linker-phosphorylated RSMAD (B and E), and (C and F) ppERK. Receptor endocytosis bias via either clathrin-coated pits (red) or lipid rafts (green) was simulated by increasing 10-fold the rate constant related to one internalization mechanism and leaving constant the corresponding value related to the other, and vice-versa. The effect of facilitated recycling of receptors from rafts was also examined by increasing the rate of receptor recycling 10-fold in addition to induced endocytosis bias (D, E, and F). Corresponding responses under nominal conditions are in blue lines for comparison.

Another question of interest concerns how the intensity of ppERK can be elevated at the same time that SMAD activation is attenuated, if, as we have shown, shifting the balance to make raft-mediated receptor trafficking predominant alone is unable to achieve this objective. The stated goal may be achievable if receptors internalized via lipid-rafts can avoid undergoing fast ligand-induced degradation. To assess the plausibility of this proposition, we increased the rate of recycling of raft receptors 10-fold in conjunction with the 10-fold increase in the rate of raft-mediated internalization investigated earlier. Figures 7D-F show that when *both* raft-mediated internalization *and* recycling of receptors are facilitated, SMAD activity is still attenuated (green in Figure 7D), but this time the intensity of ERK activity is significantly increased, compared to nominal conditions, and becomes transient (green in Figure 7F) as in the nominal case. ERK-phosphorylated SMAD becomes markedly enhanced as well (green in Figure 7E) in a slightly different but essentially similar manner as ERK. Collectively, these results imply that the induction of a specific ERK-mediated outcome depends on how fast the receptors are recycled. Specifically, facilitating raft-mediated receptor endocytosis alone (with essential no recycling) will enhance the expression of ERK-inducible genes that are primarily intensity-dependent (but not duration-dependent), while accelerated receptor recycling along with rapid raft-mediated internalization will preferentially promote the expression of duration-sensitive (and intensity-insensitive) genes.

***Taken together, these results suggest that the balance between compartment-specific receptor endocytosis plays a critical role in determining TGF-*β*-induced cellular outcomes. Specifically, clathrin-dominant receptor internalization may preferentially induce SMAD-mediated responses, while raft-dominant receptor internalization may drive various ERK-dominant outcomes, depending on the speed of receptor recycling*.**

### Simulation of Cancerous Cells Signaling Characteristics

In addition to providing the sort of general insight into the mechanisms of ligand-dependent control of SMAD and ERK activity discussed thus far, the model can also be used to generate testable predictions of TGF-β-induced behavior of complex biochemical processes— predictions that could provide new mechanistic insight. Of particular interest to us is how known, cancer-correlated network abnormalities affect the dynamic behavior of the TGF-β signaling system, and what differentiates abnormal and normal behavior. We believe that such differences between normal and cancerous system responses can offer clues into the mechanisms by which epithelial-derived cancer cells exploit TGF-β signaling for their progression and metastasis. What follows is an investigation (via carefully designed simulations) of the dynamic behavior of cancer cells in the context of TGF-β-induced SMAD and ERK signaling, based on the most common features of cancer cells reported in the literature.

Cancer, a broad collection of complex, context-dependent, dynamic diseases, is difficult to characterize with a few features. Nevertheless, there is current consensus regarding certain characteristics common to many cancer cells. For our purposes here, the most relevant are that cancer cells have the capacity to avoid or resist apoptosis mediated by the tumor suppressive effects of TGF-β [39]; and that abnormal alterations (e.g., mutation, or up/down-regulation) of the core components of the TGF-β pathways drive tumorigenesis [40]. Of all the components of the TGF-β pathways, TGF-β receptors (either type I or type II or both) are the most frequent targets of abnormal alterations (e.g., deletion, mutation, or downregulation) observed in a wide variety of primary human cancer types [40]. These aberrant alterations result directly in the reduction in the number of functional TGF-β receptors, rendering cancer cells less responsive to TGF-β stimulation [12]. In the context of MAPK signaling, the signaling component most frequently targeted for abnormal alterations is Ras protein. First, the *ras* gene is frequently mutated in human cancers: approximately 30% of human cancers express mutant Ras, termed ‘Ras oncoprotein’ [41–43]. This mutation impairs intrinsic GTPase activity of Ras and renders Ras resistant to GAP, a mediator of GTP hydrolysis for Ras inactivation. Consequently, once in the activated state, the Ras oncoprotein is unable to turn itself off and thus accumulates in the hyperactive, GTP-bound conformation. In addition, Ras oncoprotein is also overexpressed in a variety of human cancers, including neuroblastomas, esophageal, head and neck, laryngeal, thyroid, lung, liver, intestinal, gastric, colorectal, breast, bladder, endometrial, ovarian tumors and leukemias, making it an important prognostic marker [41].

Furthermore, the level of lipid rafts, cholesterol-enriched membrane microdomains, is known to be altered in cancer. A number of studies have demonstrated that cholesterol accumulates in several solid tumors, including prostate and oral cancers [44, 45], and that such accumulation can disturb cell signaling significantly [46, 17]. In particular, it is reported that elevated cholesterol levels can increase the formation and/or stabilization of lipid rafts/caveolae by integration into the plasma membrane, thereby increasing the localization of TGF-β receptors in lipid rafts/caveolae [38].

On the basis of these facts, we investigated the effect of some of these common abnormal alterations on the TGF-β signaling system, by simulating the system response to a single step of magnitude 80 pM of TGF-β while implementing the following changes on the indicated components: (i) a 10-fold reduction in the initial level and production rate of both Type I and Type II receptors (as in our previous study [12]); (ii) a 10- or 100-fold elevation of the initial level of RasGTP; (iii) a zero rate of GDP conversion; (iv) a 10-fold increase in the rate of lipid raft-mediated receptor internalization; (v) a 10-fold increase in the rate of receptor recycling from rafts. How these changes were combined to generate a collection of 9 distinct simulations is shown in Table 1.

**Table 1.** Conditions employed in simulating cancerous signaling behavior.

|  | Abnormality | Simulation | 1 | 2 | 3 | 4 | 5 | 6 | 7 | 8 | 9 |
| --- | --- | --- | --- | --- | --- | --- | --- | --- | --- | --- | --- |
| **Normal** | None | N | O |  |  |  |  |  |  |  |  |
| **Cancerous** | TGFβ receptor loss | TR | | O | O | O | O | O | O | O | O |
|  | Ras overexpression | RO1 |  |  | O |  |  |  |  | O | O |
|  |  | RO2 |  |  |  | O |  |  |  |  |  |
|  | Ras mutation | RM |  |  |  |  | O | O | O | O | O |
|  | Cholesterol perturbation | CR |  |  |  |  |  | O | O | O | O |
|  |  | CC |  |  |  |  |  |  | O |  | O |

The indicated simulation conditions are coded as follows :
N Nominal conditions
TR 10-fold decrease in the initial levels and synthesis rates of TβRI/II
RO1 10-fold increase in the Ras level
RO2 100-fold increase in the Ras level
CR 10-fold increase in the rates of raft-mediated internalization
CC 10-fold increase in the rates of receptor recycling from rafts

Thus, for example, Simulation #1 represents nominal behavior for a normal cell with no abnormalities; Simulation #2 represents the behavior when the only abnormality is a 10-fold loss in TGF-β receptor activity; and Simulation #7 involves a combination of 4 abnormalities: TGF-β receptor activity loss, Ras mutation (implemented via a 100-fold decrease in the rate of hydrolysis of RasGTP), and both versions of “cholesterol perturbation” (a 10-fold increase in *both* the rates of raft-mediated internalization and the rates of receptor recycling from rafts). The complete set of 9 simulation results is summarized in Figure 8.

**Figure 8.**
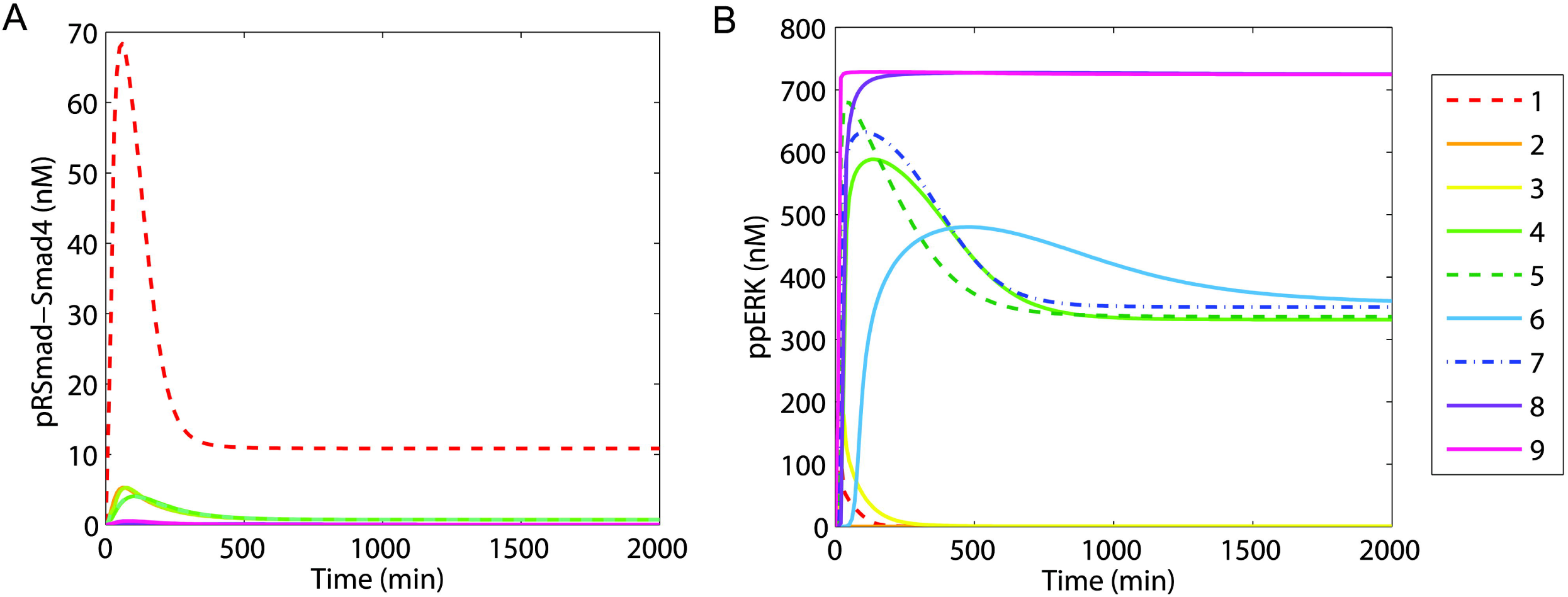
Model predictions under various cancerous conditions: (A) pRSMAD-SMAD4 complex and (B) ppERK responses to a step change in TGF-β stimulation. Numbers in the legend refer to the simulation indices in Table 1.

A direct comparison of normal versus cancerous SMAD signaling responses reveals that SMAD activation is significantly attenuated under *all* abnormal conditions (Figure 8A). As observed in our previous SMAD model, a sharp drop in the level of functional TGF-β receptors leads to a marked decrease in the activity of receptor-activated SMAD complex. From a control theory perspective, the loss of functional TGF-β receptors corresponds to a reduction in the “process gain,” as a result of which only a small portion of the input signal is “transduced” into the response [47, 48]. If, in addition to the loss of receptors, receptor internalization is biased toward lipid rafts, then activity of the SMAD complex is reduced even further, to an almost undetectable level. However, once the receptor levels are lowered significantly, other ERK-signaling-associated abnormalities will no longer lead to any noticeable change in the dynamics of activated SMAD complex. Consequently, the implication is that the activity of receptor-activated SMAD complex depends mostly on the dynamics and the levels of functional receptors. Figure 8A shows this clearly, where, beyond the sharp decrease in SMAD complex activity captured in Simulation #2 (TGF-β receptor loss), conceptually, there is not much to distinguish one response from another.

The situation with ppERK in Figure 8B is entirely different. While the loss of TGF-β receptors alone can result in a sharp decrease in the level of ppERK almost to the point of disappearing altogether (compare Simulation #1 to the barely visible response #2)—indicating that the activity of ppERK is also affected significantly by the level of functional receptors— depending on the type and degree of abnormalities in question, the 9 responses differ considerably. Figure 8B is therefore significantly more informative.

Leaving out Simulation #2 which shows a virtually non-responsive ppERK as a result of TGF-β receptor loss, the other responses may be categorized into three distinct groups:

I. Simulations #1 and #3: characterized by a transient rise to a peak value followed by exponential-like decay to zero. The conditions associated with this group are: #1, Nominal conditions; #3, Receptor loss *plus* Ras overexpression.
II. Simulations #4, #5, #6 and #7: characterized by a fairly rapid rise to a peak value followed by a steady decline to a moderately elevated but sustained value at steady state. Associated conditions: #4, Receptor loss *plus* severe Ras overexpression; #5, Receptor loss *plus* Ras Mutation; #6, same as #5 *plus* a 10-fold increase in the rates of raft-mediated internalization; #7, same as #6 *plus* 10-fold increase in the rates of receptor recycling from rafts.
III. Simulations #8 and #9: characterized by a sharp, swift rise that is sustained at the highest level at steady state, neither declining nor increasing. These two responses are differentiated only by the speed with which steady state is attained, which is only a bit slower in #8, otherwise both settle down to the same steady state high value. Associated conditions: the entire complement of abnormalities in #9, with only receptor recycling from rafts missing in #8.

Group I responses indicate that a combination of Ras overexpression and TGF-β receptor loss results in ppERK dynamics similar in characteristics to nominal behavior, differing only in terms of magnitude and speed of response. These responses, when compared with those in Group II, show that to obtain significantly elevated and prolonged ERK activation requires at the very least, activating Ras mutation (#5, #6, #7), or extreme Ras overexpression (#4). We may also observe by comparing Simulations #4 and #5 that Ras mutation (impairing its GTPase activity) speeds up the elevation of ppERK activity more than Ras overexpression.

The simulations in Groups II and III demonstrate the influence of receptor internalization and recycling on ERK activity. The response in Simulation #6, with its distinctive delay and sluggishness, differs somewhat from the other responses in the same Group II. Such a delay in ERK activation is directly attributable to unbalanced receptor internalization via lipid rafts; and the fact that this effect can be partially offset by rapid recycling of raft receptors is responsible for the more rapid response in Simulation #7, which is similar to Simulation #5 where there are no abnormalities related to receptor internalization or recycling. Note that these observations are consistent with Figures 7C and 7F. The addition of Ras overexpression to the conditions in Simulation #6 produces the response in Simulation #8, with a much shorter delay and a much faster approach to the steady state sustained and elevated value of ERK activity. The further addition of a 10-fold increase in the rates of raft-mediated receptor recycling results in Simulation #9, which similar to Simulation #8, but much faster, and ultimately sustained at the same hyper-activated steady state level.

Collectively, these *in silico* experiments suggest that in response to TGF-β stimulation, the dynamic characteristics of both SMAD and ERK signaling change significantly during cancer development: ***Normal cells exhibit enhanced and sustained SMAD signaling with transient ERK activation, whereas cancer cells produce elevated and prolonged ERK signaling with enervated SMAD signaling.*** These distinctive differences between normal and cancerous signaling behavior provide clues regarding what might be responsible for the apparently contradictory roles played by TGF-β during cancer development, as we discuss next.

### A Hypothesis on the Dual Role of TGF-β in Cancer

The results in the previous section suggest that as cancer advances, the effect of SMAD signaling becomes less significant, almost to the point of total lack of response, while the effect of ERK signaling becomes more pronounced. This observation leads to the following hypothesis regarding TGF-β’s paradoxical roles in cancer where the cytokine, a potent tumor-suppressor in normal cells, appears to become a tumor promoter in cancerous cells.

*Sustained and/or enhanced SMAD activation is required to induce tumor suppression while sustained and/or enhanced ERK activity is required for tumor promotion. In normal cells, TGF-* β *stimulation produces primarily sustained and sufficiently high ligand-induced SMAD activation, in conjunction with transient and relatively low ERK activity; the former is sufficient to induce transcription of tumor suppressor genes, while the latter is not strong enough for tumor promotion. In cancer cells, TGF-*β *stimulation produces enhanced and prolonged TGF-*β*-induced ppERK activity, in conjunction with significantly reduced and potentially insufficient SMAD activation; the former is sufficient to induce pro-oncogenic responses, while the latter is too weak to induce tumor suppression*.

Consequently, we observe that for normal cells, the net cellular readout in response to TGF-β stimulation will be tumor suppressive (see left-hand side of Figure 9); on the other hand, the net phenotypic response in cancer cells will be tumor-promoting (right-hand side Figure 9).

**Figure 9.**
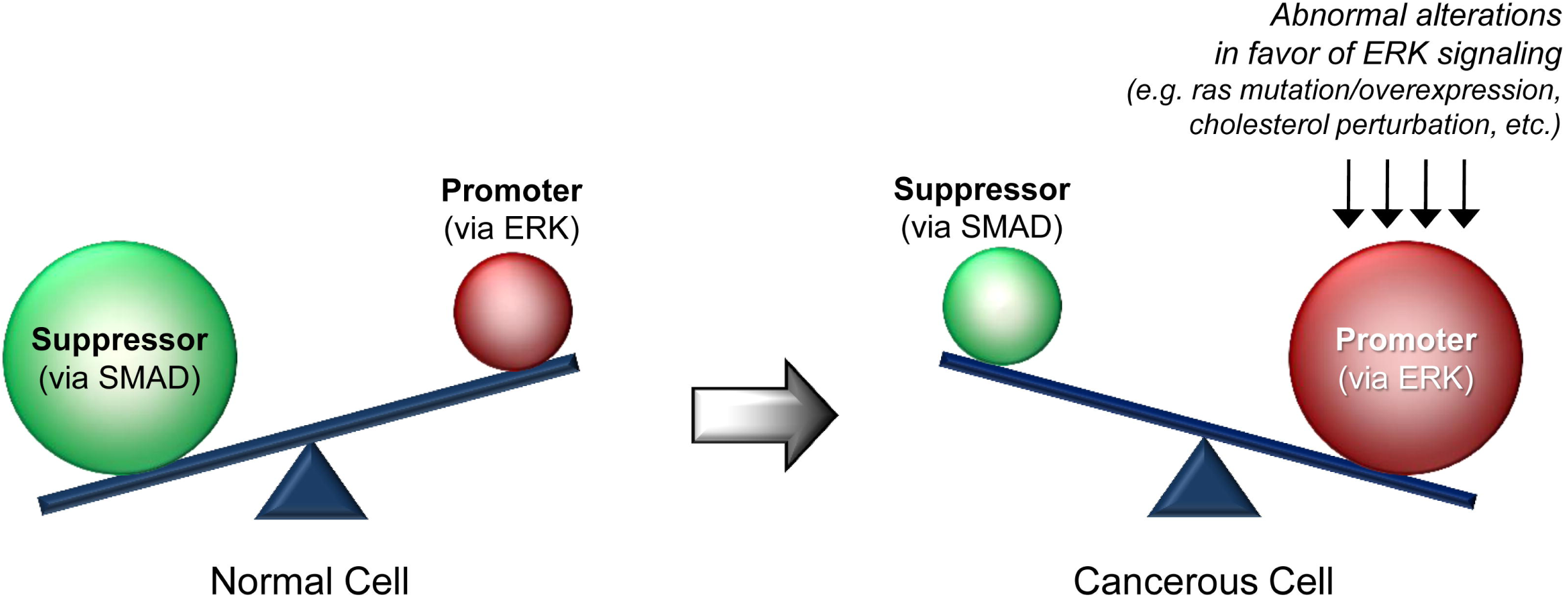
Schematic representation of a postulated hypothesis regarding the dual role of TGF-β during cancer progression.

Indeed, this hypothesis and its consequences are supported in part by previous experimental results that show that sustained activation of the ERK1/2 pathway contributes to oncogenic transformation or cell migration [49–51]. In particular, mounting evidence indicates that the duration of ERK signaling influences cellular outcomes. For example, it was reported that sustained Raf-MEK-ERK activated by oncogenic Ras is required for induction of β3 integrin transcription (which is associated with particularly elevated tumor aggressiveness), whereas transient activation of Raf-MEK-ERK signaling by growth factors and mitogens had no such effect [52]. Similarly, it has been suggested that sustained, but not transient, activation of ERK is necessary for inducing S-phase entry by down-regulating anti-proliferative genes until the onset of S-phase to allow successful G1 phase progression [53]. This means that while SMAD-mediated anti-tumorigenic activity such as induction of cell cycle arrest or apoptosis are attenuated, enhanced and prolonged ERK signaling may initiate the cell cycle progress or other tumor promoting mechanisms depending on cellular context.

To summarize, our hypothesis is that abnormal alterations in the molecular context, which promotes changes in the intensity and duration of SMAD and ERK signaling during cancer progression and which ultimately leads to a tipping of the balance between the SMAD and ERK pathways in favor of tumor promotion, may be responsible for the two apparently opposing roles that TGF-β plays in cancer development. In fact, the basic concept of our hypothesis is consistent with other hypotheses postulated previously in the literature, but without quantitative supporting evidence: the imbalances between SMAD and alternative signal transduction pathway(s) may underlie the ability of TGF-β to induce pro-oncogenic responses such as EMT [54, 55]. Our study is the first *quantitative* description of this hypothesis, which has until now been described only qualitatively.

### Therapeutic Implications

As a result of the discussion in the above section, we may now observe that the fundamental challenge of any TGF-β-based cancer therapeutic is two-fold: restore lost tumor suppressor function, *and* simultaneously inhibit late-developing pro-oncogenic effects [54]. Previously, we proposed as a promising therapeutic option, the re-sensitization of cancer cells to the anti-tumor function of TGF-β [12, 47], achievable *in principle* via the restoration of normal TGF-β receptor expression to a healthy level. However, the simulation results shown in Figure 8 suggest that for some cancer cells, this alone may not be enough. This is because even with a restored SMAD pathway, oncogenic Ras can still interfere with TGF-β-induced signaling, so that ppERK activity may still remain dangerously elevated. As such, while the re-sensitized SMAD pathway may restore tumor suppression action, the pro-oncogenic override action effected via the ERK pathway remains.

To establish this point quantitatively, we simulated the effect of receptor recovery (achieved by restoring the expression of TGF-β receptors to normal levels) under a single cancerous condition: Ras protein mutation leading to loss of GTPase activity. The results in Figure 10 show the system response to the same single step input in TGF-β used to generate Figure 8, under nominal conditions (no abnormalities: blue), with the dual abnormalities of TGF-β receptor loss and Ras mutation (red), and with receptor recovery and unchanged Ras mutation (green). Observe that with receptor recovery, SMAD response improves significantly (Fig. 10A), especially at steady state where the original sustained value is attained. However, peak activity of SMAD complex is significantly lower than under nominal conditions. On the contrary, the effect of receptor recovery on either the intensity or the duration of ppERK activity is essentially inconsequential (Fig. 10C), especially at steady state. Furthermore, the elevated and sustained ERK activation occurring under receptor recovery leads to enhanced phosphorylation of the SMAD linker regions (Fig. 10B), which in turn prevents SMAD-mediated transcription in the nucleus. The clear implication is that even when TGF-β receptor levels are restored, cancer cells may not be responsive fully to the anti-growth effects of TGF-β, but will definitely remain susceptible to the acquired pro-oncogenic effect because of the still amplified and prolonged activity of ppERK. Consequently, to be effective, a targeted therapy regimen for treating a cancer patient with a genetic signature of oncogenic Ras requires *at least* a combination therapy to restore *both* the expression of TGF-β receptors *and* the GTPase activity of Ras. It is possible that restoration of the canonical SMAD pathway will make cancer cells more sensitive to chemotherapy induced apoptosis [56, 57]. Clearly, such insight is not possible from a narrow focus on the SMAD pathway alone. By the same token, however, we must also caution that if there are other pathways that interfere significantly with TGF-β-induced SMAD and ERK signaling, the results of this study must be appropriately modified to account for those interactions as well. But this is precisely the main point of this study: that in developing quantitative understanding and subsequent targeted cancer therapy, a signaling *network* should be modeled as comprehensively as possible, even if the comprehensive network model is developed sequentially, adding more components and increasing model complexity successively as necessary.

**Figure 10.**
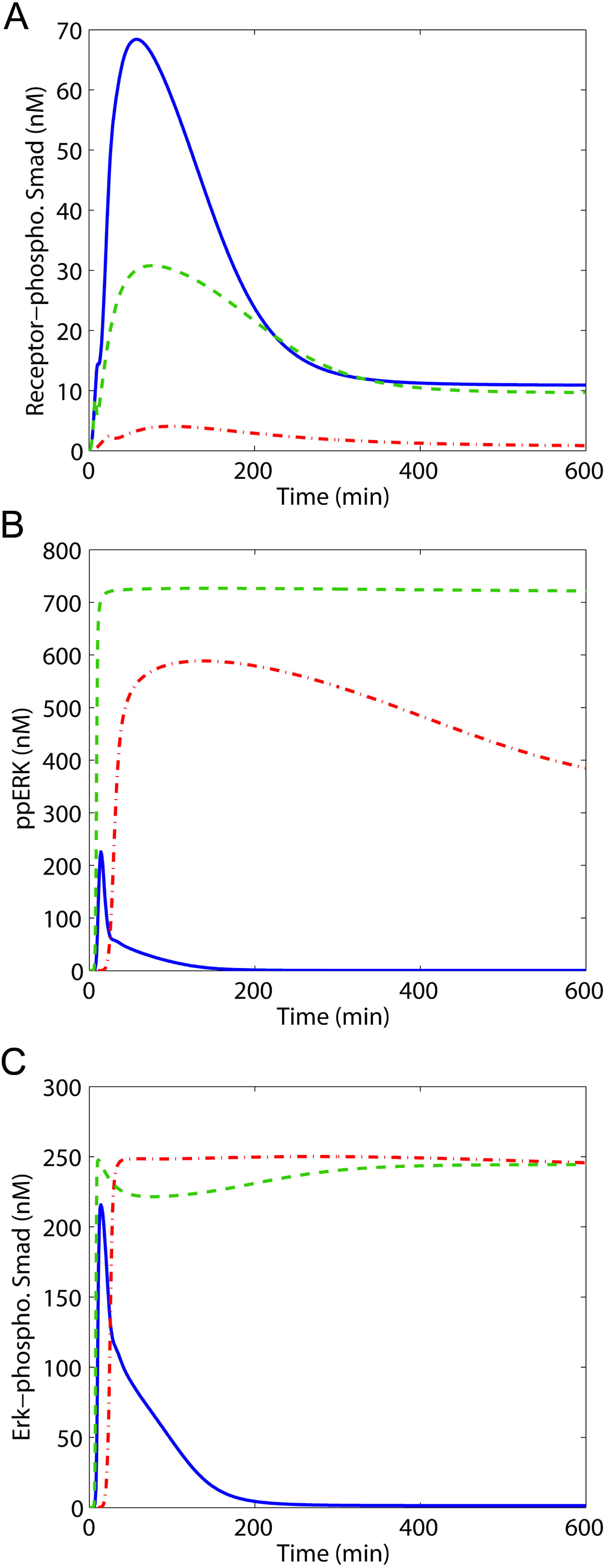
Effect of restoration of TGF-β receptor functionality in cancerous cells. Model prediction of the dynamic responses of (A) pRSMAD-SMAD4 complex, (B) ppERK and (C) SMAD phosphorylated in the linker region, to sustained TGF-β stimulus; under nominal conditions (blue), with dual abnormalities of receptor loss and Ras mutation (red), and with receptor recovery and unchanged Ras mutation (green).

## Conclusions

Mathematical modeling of signal transduction pathways continues to provide *quantitative* insight into how extracellular signals are communicated and integrated intracellularly in order to control cellular responses. As a result of the intrinsic complexity associated with each pathway, pioneering modeling efforts were, understandably, restricted to single, well-characterized pathways studied in isolation. Of course, it is inconceivable that *all* signal transduction pathways act in isolation, without some degree of mutual interactions whereby primary signals within one pathway influence and are influenced by those within other pathways. Consequently, mathematical models of isolated signal transduction pathways must be recognized properly as *first order approximations* of potentially more complex characteristics.

Nevertheless, the mere presence of crosstalk does not necessarily mean that the implied interactions and cross-influences are all uniformly strong or consequential. In the case of TGF-β signaling, we have established that the activity of the canonical SMAD pathway is significantly affected by signaling through other pathways, especially the ERK pathway. By incorporating such crosstalk into our previously published canonical SMAD pathway model, we have been able to provide additional insight not possible otherwise. As one might expect, however, beyond providing insight, the presence of significant crosstalk has many implications, especially for the design of targeted therapeutics that are based on remedying signal transduction malfunction. Must one then have to model the complete collection of signaling circuits within the cell, in addition to the full complement and exploit such understanding for practical applications? We believe that the answer must be *no*. For one thing, not all interactions are significant; for another, even if it were somehow possible to develop such a fully comprehensive model, the end result would be too complex, contain too many unidentifiable parameters, and be far too unwieldy to be useful. In chemical process control, virtually all processes of industrial significance are multivariable, where to a greater or lesser extent, all manipulated variables influence all controlled variables; and yet single loop control systems are still prevalent and many still function effectively. Not all chemical processes require full multivariable controllers for effective operation [48].

An important perspective of this study may therefore be stated as follows: to develop meaningful and useful quantitative understanding of a signaling *network* requires as comprehensive a model as possible, but the complexity of biological signaling systems and the necessity for usefulness of the resulting model may require that the comprehensive network model be developed sequentially. We believe that useful models of signal transduction pathways can (and maybe *should*) be developed by starting with an eye toward producing the highest possible fidelity with the simplest possible model, only adding more components and increasing model complexity successively as necessary.

## Author Contributions

Conceived and designed the experiments: SWC and BAO. Performed the experiments: SWC. Analyzed the data: SWC and BAO. Provided experimental background: CRC, MCFC. Wrote and reviewed the paper: SWC, BAO, CRC, MCFC.

## Additional Files

Additional file 1. Supplementary tables: Table S1 - Model equations; Table S2 - Initial concentrations; Table S3 - Model parameters.

Additional file 2. Figure S1. Schematic representation of the TGF-β-induced SMAD and ERK signaling pathways.

## Acknowledgments

This work was supported by the Department of Defense grant PC050554, and by the Institute for Multiscale Modeling of Biological Interactions (IMMBI) (funded by the Department of Energy). Additional work was supported by National Institutes of Health/National Cancer Institute P01 CA098912, the University of Delaware Research Foundation, and the National Institutes of Health INBRE P20RR016472.

