## Supplementary figures and images for "Computational Modeling and Analysis of the TGF-β-induced ERK and SMAD Pathways"

### Supplemental figure 1

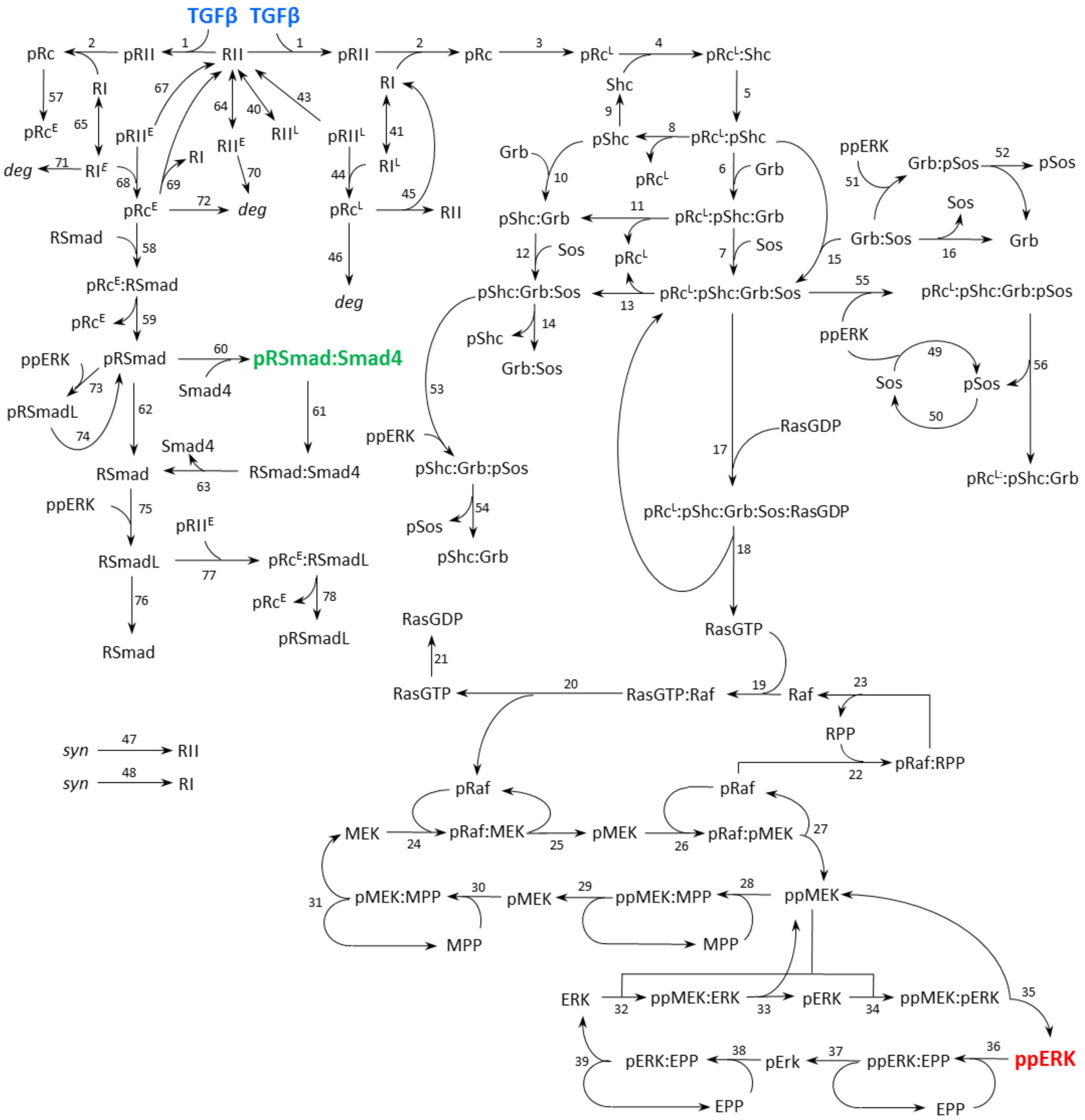
